# Adaptive, arousal-related adjustments of perceptual biases optimize perception in a dynamic environment

**DOI:** 10.1101/083766

**Authors:** Kamesh Krishnamurthy, Matthew R. Nassar, Shilpa Sarode, Joshua I Gold

**Author notes:** contributed equally and listed alphabetically (Kamesh Krishnamurthy, Matthew R. Nassar). Corresponding Author: Joshua I. Gold, Department of Neuroscience, University of Pennsylvania, Philadelphia, PA 19104-6074.

## Abstract

Prior expectations can be used to improve perceptual judgments about ambiguous stimuli. However, little is known about if and how these improvements are maintained in dynamic environments in which the quality of appropriate priors changes from one stimulus to the next. Using a novel sound-localization task, we show that changes in stimulus predictability lead to arousal-mediated adjustments in the magnitude of prior-driven biases that optimize perceptual judgments about each stimulus. These adjustments depend on task-dependent changes in the relevance and reliability of prior expectations, which subjects update using both normative and idiosyncratic principles. The resulting variations in biases across task conditions and individuals are reflected in modulations of pupil diameter, such that larger stimulus-evoked pupil responses correspond to smaller biases. These results suggest a critical role for the arousal system in adjusting the strength of perceptual biases with respect to inferred environmental dynamics to optimize perceptual judgements.

## Introduction

Perception is shaped by prior expectations (“priors”) on the statistical structure of the sensory world (Bar, 2004; Edwards, 1965; Link and Heath, 1975; Maddox and Bohil, 1998; Seriès, 2013; Summerfield and Egner, 2009). When the environmental statistics are stationary and well known, priors on those statistics can bias the perception of relevant sensory stimuli (Fischer and Peña, 2011; Knill and Pouget, 2004). For example, the prevalence of relatively slow-versus fast-moving objects in the world can lead to biases in the perception of object speed (Stocker and Simoncelli, 2006). However, many environmental statistics that are relevant to perception can be highly non-stationary. For example, the locations of sources of sensory input are constantly changing relative to a given observer. The goal of this study was to examine how priors on such dynamic features of the environment are updated and used to shape perception.

To achieve this goal, we developed a novel auditory-localization task that required human subjects to both predict and report the perceived location of a simulated sound source as the predictability of the location varied over time (Fig. 1a–c). The statistical structure of the task is similar to ones we used previously to show that people can make effective predictions in dynamic environments by adaptively modulating the influence of new information on existing beliefs (Nassar et al., 2010; Nassar et al., 2012). However, unlike in those tasks, here we used perceptually ambiguous stimuli that allowed us to focus on a novel question: how do dynamic fluctuations in the ability to make effective predictions affect the magnitude of perceptual biases towards expected stimulus values? In principle, these perceptual biases could be adjusted through a form of optimal (“Bayesian”) inference that takes into account dynamic changes in the priors (Knill and Richards, 1996; Nassar et al., 2010; Wilson et al., 2010). Specifically, as long as the statistical structure of the sampled locations in our task remain stable, new sounds can be used to develop increasingly reliable priors about the locations of subsequent sounds. These increasingly reliable priors should, in turn, have an increasingly strong and beneficial influence on the perception of those sounds, reducing localization errors (Fig. 1d,e). However, the statistics of the sampled locations can undergo abrupt change-points that render previously held priors irrelevant to new sounds. These seemingly reliable but irrelevant priors should not influence the perception of sound-source location, which under these conditions should be limited entirely by sensory uncertainty (Fig. 1f).

**Figure 1:**
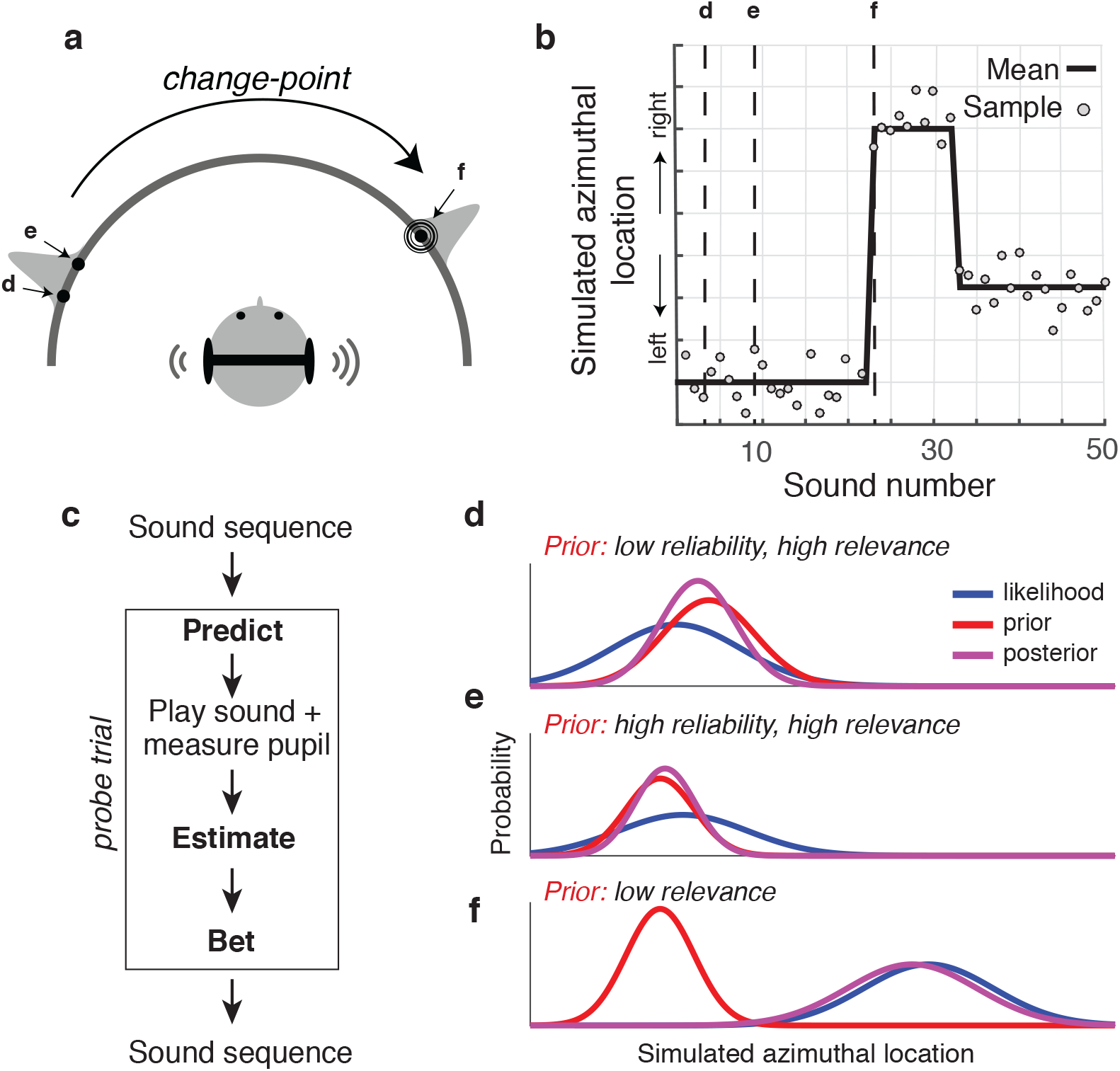
Dynamic sound-localization task. (**a**) Subjects listened via headphones to noise bursts with virtual source locations that varied along the frontal, azimuthal plane. The locations were sampled (labeled arrows) from a Gaussian distribution (gray) with a mean that changed abruptly on unsignaled change-points (probability=0.15 for each sound) and a STD of 10° in low-noise blocks, 20° in high-noise blocks. The subjects listened passively to the sound sequence, except for occasional probe trials. All sounds except the probe sound were presented simultaneously with their corresponding locations on a semicircular arc shown on the isoluminant visual display, allowing subjects to develop priors on sound-source location based on both the auditory and visual signals and maintain a stable mapping between the two. (**b**) An example trial sequence showing the mean (solid line) and sampled (points) locations over 50 trials. Vertical dashed lines indicate randomly selected probe trials. (**c**) Probe-trial sequence. Using a mouse to control a cursor on the visual display, the subject reported: 1) the predicted location of the upcoming probe sound, followed by 250-ms fixation, presentation of the probe sound, then continued fixation for 2.5 s to allow for pupil measurements; 2) the estimated location of the probe sound; and 3) a high or low bet that the true location was within a small window centered on their estimate. The sound sequence then continued until the next probe. (**d–f**) Schematic illustrating the changing reliability and relevance of priors for the probe sounds in **a** and **b**, as indicated. Given a fixed-width likelihood function, more reliable and relevant priors have a stronger and more beneficial influence on the percept, here represented as the posterior, which is least uncertain (narrowest) in **e** and most uncertain in **f**.

We also measured pupil diameter, an index of arousal that is thought to reflect the activation of the locus coeruleus (LC)-norepinephrine (NE) system, while subjects performed the task (Aston-Jones and Cohen, 2005; Joshi et al., 2016). Pupil diameter tracks the extent to which predictions are updated in response to new information in dynamic and perceptually unambiguous cognitive tasks (Nassar et al., 2012). Here we tested the hypothesis that such changes in arousal play an important role in shaping perception. In particular, we examined whether the arousal system controls the extent to which perceptual judgments about ambiguous sensory stimuli are biased toward prior expectations in accordance with the relevance and reliability of those expectations.

Our results yield new insights into the relationship between perception and arousal. We show that the subjects’ priors had a variable influence on their perceptual reports. This variability was predicted by changes in the relevance and reliability of those priors, across task conditions and individual subjects. These effects were encoded in both baseline and stimulus-evoked changes in pupil diameter, such that larger diameters corresponded to less influence of priors on the perception of that stimulus. Taken together, these findings support a fundamental role for pupil-linked arousal systems, including the LC-NE system, in adaptively adjusting the influence of priors on perception in accordance with environmental dynamics.

## Results

Twenty-nine subjects performed both the dynamic localization task (Fig. 1) and a control task that required perceptual reports of simulated sound-source locations that lacked predictable, sequential structure. Overall, the subjects tended to perform both tasks in an effective manner, providing predictions on the dynamic task and perceptual reports on both tasks that corresponded strongly to the simulated sound-source locations (Fig. 2). On the control task, the Pearson’s correlation between simulated and reported location had median [interquartile range, or IQR] values of 0.926 [0.895–0.944] across subjects (Fig. 2a,d). On the dynamic task, there were similarly high correlations for both the predictions and perceptual reports (predictions on non-change-point trials: *r*=0.907 [0.895–0.921], Fig. 2b,e; perceptual reports on all trials: *r*=0.948 [0.941–0.964], Fig. 2c,f). However, the subjects also tended to make errors that varied considerably from trial to trial on both tasks (Fig. 2g–i). Subsequent analyses focus on how the subjects minimized their errors on the dynamic task by exploiting the fluctuating predictability of sound-source locations on that task.

**Figure 2:**
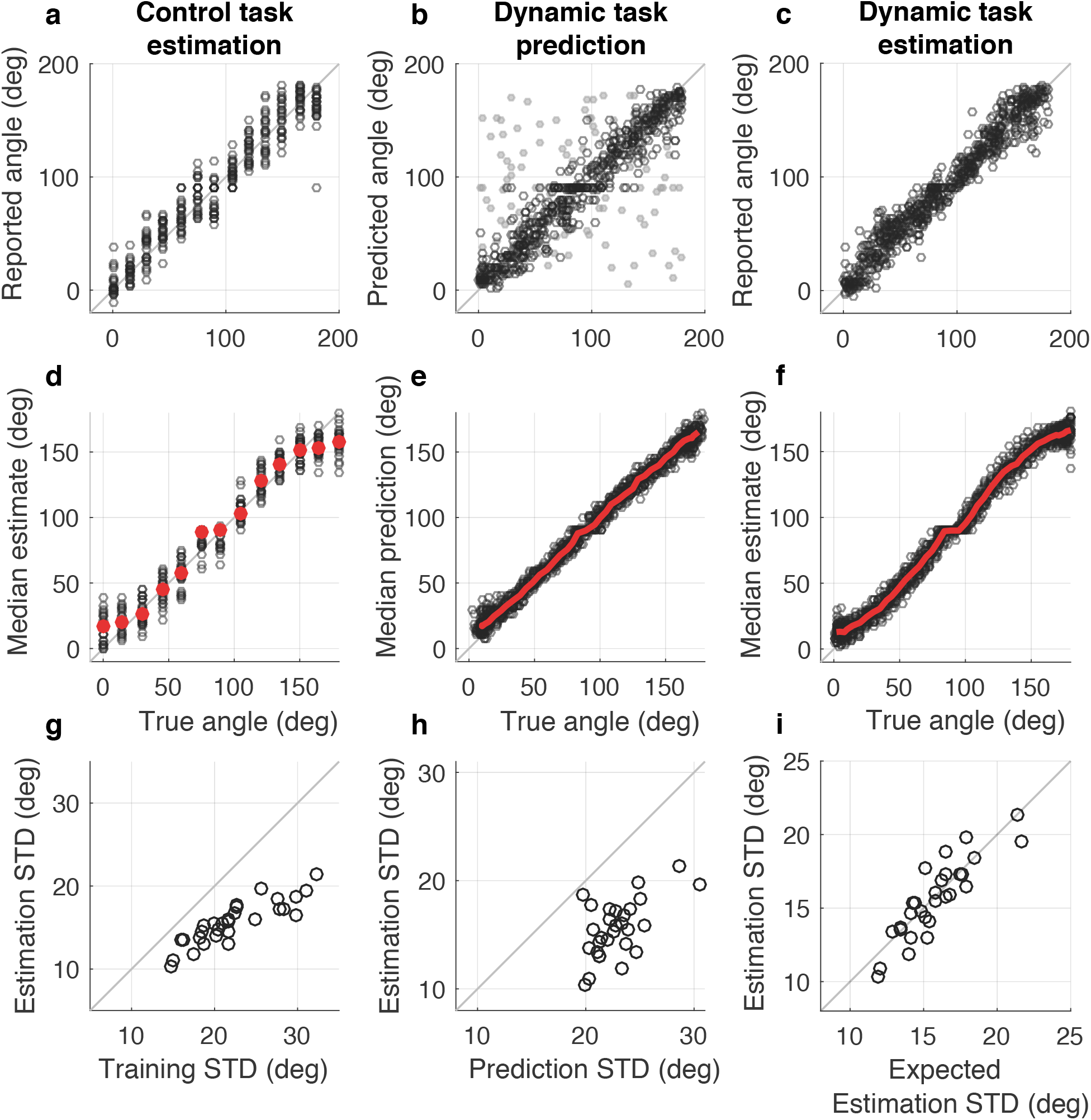
Overall prediction and estimation performance. (**a**–**c**) Reported versus true (simulated) sound-source angle for an example subject for: (**a**) estimations from the control task; (**b**) predictions from the dynamic task (light gray points indicate change-point trials, on which the probe location was, by design, unpredictable); and (**c**) estimations from the dynamic task, including all trials. (**d**–**f**) Population summaries, plotted as in (**a**–**c**), with per-subject median values shown in black and the median of medians shown in red. For the dynamic tasks, median values were calculated in sliding 20° windows. Non-change-point trials were excluded from the predictions in (**e**). Note that the subjects’ perceptual reports (**d** and **f**) were biased slightly towards straight ahead at the far periphery. This bias, which likely reflected learned expectations that sounds were only played in the frontal plane, is accounted for in later analyses (β_5_ and β_6_ in Eq. 5). (**g**–**i**) STD of the perceptual errors from the dynamic task plotted versus the STD of: (**g**) the perceptual errors from the control task; (**h**) the prediction errors from the dynamic task; or (**i**) the expected STD of the perceptual errors, computed from the optimal, reliability-weighted combination of the control perceptual errors and the dynamic prediction errors. Points in **g**–**i** represent data from individual subjects. Prediction and perceptual errors were computed with respect to the simulated location of the probe sound.

### Dynamic, task-dependent modulation of perceptual biases

The subjects used both sensory and prior information to guide their perceptual reports on the dynamic task. We measured performance in terms of the variability of the distribution of trial-by-trial errors (quantified as the standard deviation, or STD, and denoted as σ). This variability was lower for perceptual reports on the dynamic task than for either: 1) predictions from that task (*σ_prior_*; Fig. 2h), or 2) perceptual reports on the control task that lacked sequential predictability and thus reflected more purely sensory processing (*σ_sensory_*; Fig. 2g). Moreover, for individual subjects, these different measures of variability were related to each other, such that perceptual errors from the dynamic task were well approximated using the optimal, reliability-weighted combination of prior and sensory information (*σ^-2^_optimal_* = *σ^-2^_prior_* + *σ^-2^_sensory_*; Fig. 2i). This result implies that, on average, the subjects tended to not only use these two sources of information, but also combine them according to their relative reliabilities to optimize perceptual performance on the dynamic task.

This integration of prior and sensory information took into account the changes in the relevance and reliability of the priors that occurred throughout the dynamic task. These changes are illustrated in Fig. 3a, which shows prediction-error STDs averaged across subjects as a function of the number of sounds after a change-point, or SAC, separately for the two noise conditions. Figure 3b shows linear contrasts that captured the salient, dynamic aspects of these changes for each subject (see inset in Fig. 3e describing CP, Exp, and Noise contrasts). Specifically, on change-point trials, predictions were irrelevant and hence most variable with respect to the subsequent sound-source location (signed-rank test for *H_0_*: the median of the distribution of per-subject CP contrasts, which compared change-points to other trials=0, *p*<10^−5^). After change-points, predictions became steadily more reliable as the number of sound sources experienced from the new distribution increased in both noise conditions (*p*< 10^−4^ for Exp_low_ and Exp_high_ contrasts, which identified linear trends across SAC 2–6 for each of the two noise conditions). The predictions were also more reliable overall in the low-versus high-noise condition (Noise contrast, *p*< 10^−5^). These dynamic trends were consistent with predictions from a normative model of predictive inference that had full knowledge of the generative statistics (Nassar et al., 2010). The model, which produced simulated predictions that were analyzed in the same way as the data, had task-dependent effects that were in the same directions and of roughly the same magnitude as the data, although the subjects tended to produce more variable predictions than the model (Fig. 3a,b diamonds).

**Figure 3:**
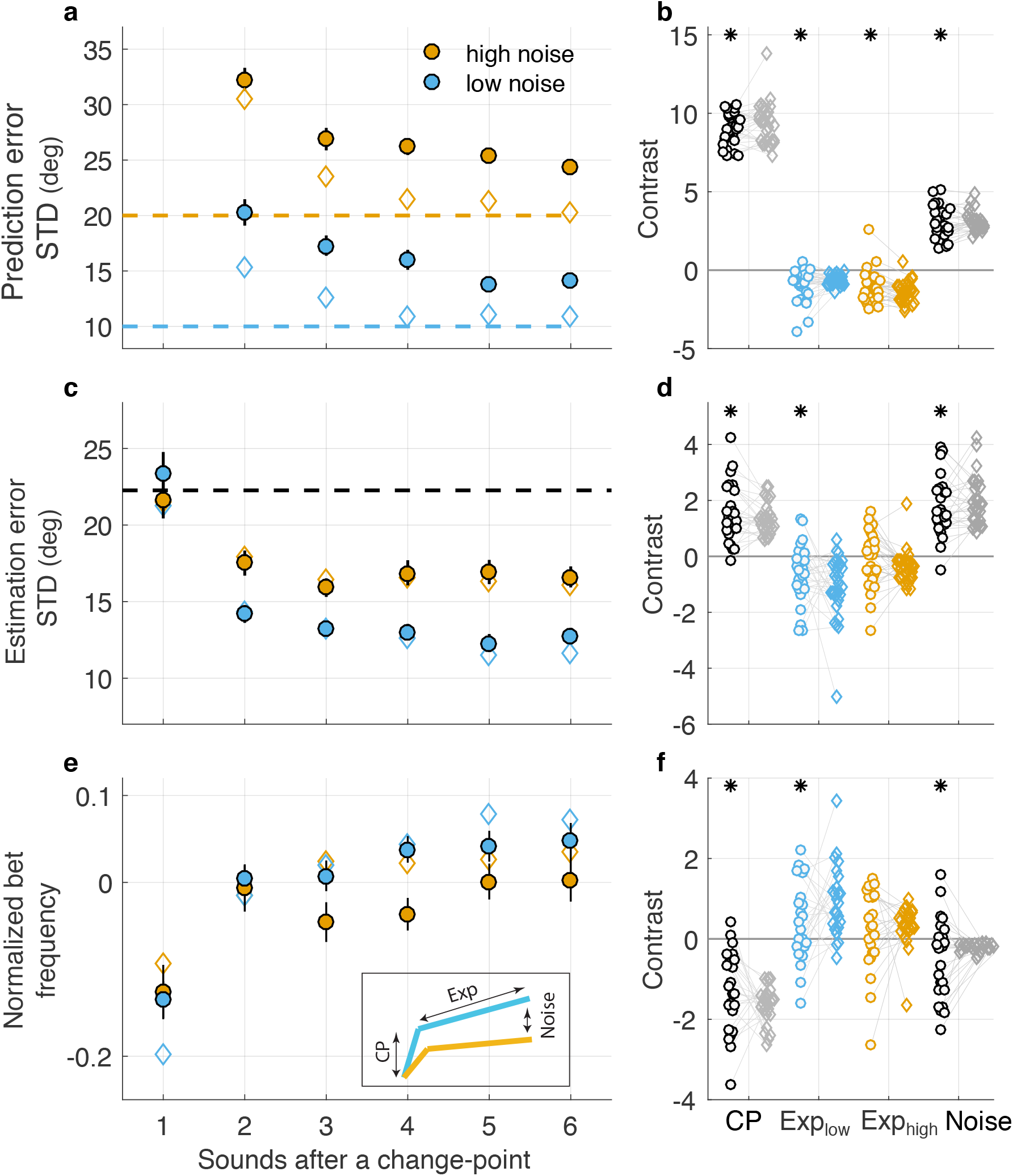
Effects of task dynamics on performance. (**a**) STD of the subjects’ prediction errors (filled circles) as a function of the number of sounds after a change-point (SAC) in the generative mean azimuthal location, plotted separately for the two noise conditions (colors, as indicated; generative STDs are shown as dashed lines). For comparison, prediction-error STDs are shown for an approximately optimal predictive-inference model (open diamonds). Data from change-point trials (SAC=1) are not shown because locations were, by design, unpredictable on those trials. (**b**) Contrast values from a linear model describing individual subject (circles) and the approximately optimal model (each diamond represents analyses based on the same sound sequence experienced by the subject connected by a line) prediction-error STD in terms of (see inset in **e**): 1) the difference between change-point and non-change-point trials (CP), 2,3) the linear trend from SAC 2–6 for low- (Exp_low_) or high- (Exp_high_) noise trials, and 4) the difference between the two noise conditions (Noise). (**c,d**) Same conventions as in **a**,**b** but for perceptual errors on the dynamic task. Diamonds represent the theoretically predicted STD of perceptual errors computed from the optimal, precision-weighted combination of the subject- and condition-specific STDs of prior errors (circles in **a**, determined separately for each subject) and the subject-specific estimation-error STDs from the control task (the median value is shown as a horizontal dashed line; see Fig. 2g). (**e,f**) Same conventions as in **a**,**b** but for the frequency of high bets relative to overall betting frequency per subject. Diamonds represent the betting frequency corresponding to the theoretical perceptual errors in **c**, computed from the fraction of the theoretical posterior distribution within the betting window. In **a**,**c**,**e**, circles and error bars are mean±sem of values measured from all 29 subjects. In **b**,**d**,**f**, points are data from individual subjects. Asterisks indicate sign-rank test for *H0*: median value from the subject data=0, *p*<0.05. In all panels, only data from sequences following noticeable change-points (changes in mean of at least twice the generative STD for SAC=1) were included.

These task-dependent changes in the subjects’ predictions were associated with similar changes in the variability of their perceptual reports (Fig. 3c,d) and their confidence in those reports, as assessed by bet frequencies (Fig. 3e,f). Perceptual-error variability tended to be higher for change-point trials, when predictions were irrelevant (CP contrast, *p*<10^−5^), and for the high-versus low-noise condition (Noise contrast, *p*<10^−5^). Perceptual-error variability also tended to decrease on experiencing more samples from the new distribution, with a reliable effect across individuals in the low-noise condition (Exp_low_ contrast, *p*<0.005) but not the high-noise condition (Exp_high_ contrast, *p*=0.4). These dynamics were also apparent in the subjects’ betting trends (Fig. 3e,f), which reflected trial-by-trial awareness of the changes in perceptual variability and included similar dependencies on CP (*p*<10^−4^), Noise (*p*=0.032), and Exp_low_ (*p*=0.03) and less reliable dependencies on Exp_high_ (*p*=0.07). Both the perceptual and betting effects were qualitatively similar, in direction and magnitude, to theoretical values computed from optimal combinations of each subjects’ changing priors (circles in Fig. 3a,b) and their fixed sensory reliability estimated from the control task (Fig. 2g; see also Fig. 1d–f). These theoretical values also showed strong effects of CP, Noise, and Exp_low_, and smaller effects of Exp_high_ (Fig. 3c–f, diamonds).

These behavioral dynamics reflected changes in the degree to which the subjects’ priors biased their perceptual reports. We quantified perceptual bias as the slope of the relationship between the prediction error and the perceptual error measured on individual trials (Fig. 4a–c). A slope of zero implies no relationship between the prediction error and the perceptual error, and thus no bias towards the prior. In contrast, slope values that increase towards unity imply increasing biases of the perceptual reports towards the prior. This perceptual bias varied systematically as a function of task conditions. Specifically, perceptual bias was lower on change-points (CP contrast, *p*<10^−5^) and for the high-versus low-noise condition (Noise contrast, *p*=0.008). Perceptual bias also tended to increase on experiencing more samples, although these effects were variable across individuals and not statistically reliable in the low-noise condition (Exp_low_ contrast, *p*=0.1; Exp_high_ contrast, *p*=0.004). These task-dependent changes in the biases were comparable in direction and magnitude to theoretically computed values given an optimal, reliability-weighted combination of the task-specific predictions on the dynamic task (circles in Fig. 3a) and fixed sensory reliability estimated from the control task (Fig. 2g), computed separately for each subject (diamonds in Fig. 4d,e). Despite these comparable task-dependent trends (compare circles and diamonds in Fig. 4e), the subjects’ perceptual biases were on average smaller than the theoretical values (compare circles and diamonds in Fig. 4d). However, overall performance, measured as perceptual-error variability, was relatively insensitive to this overall shift, as compared to task-dependent adjustments, in the magnitude of the perceptual biases (compare circles and triangles in Fig. 3c,d).

**Figure 4:**
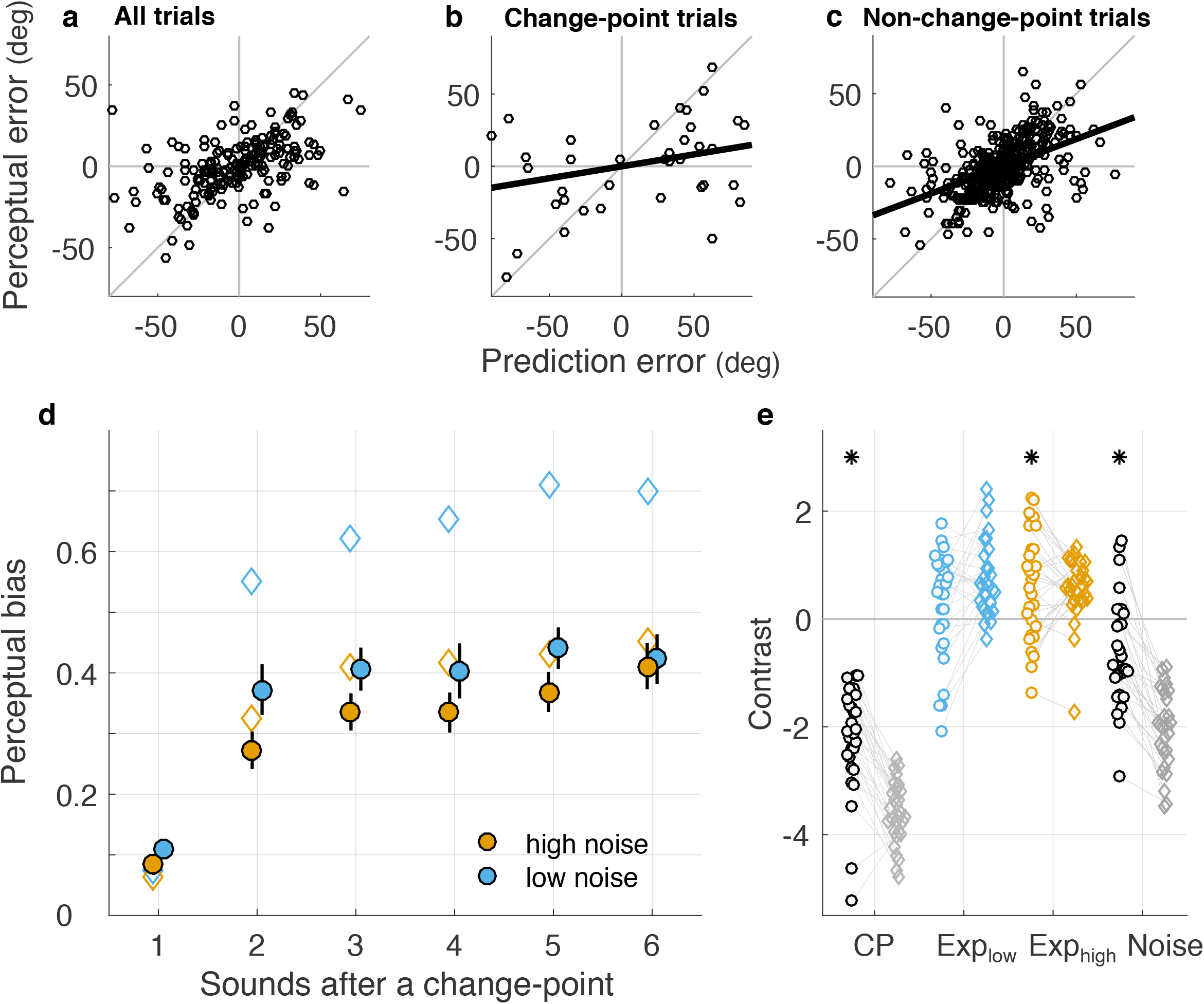
Effects of task dynamics on perceptual bias. (**a**–**c**) Example data from a single subject illustrating the quantification of perceptual bias as the slope of the best-fit line to a scatter of the perceptual error versus the prediction error. Slopes close to zero reflect a low perceptual bias (i.e., the percept is unrelated to the prediction), as on change-point trials (**b**). Slopes closer to unity reflect a higher perceptual bias (i.e., the percept more closely matches the prediction), as on non-change-point trials (**c**). (**d**) Perceptual bias as a function of the number sounds after a change-point (SAC) in the generative mean azimuthal location, plotted separately for the two noise conditions (colors, as indicated). Circles and error bars are mean±sem of values measured from all 29 subjects. Diamonds indicate the theoretically predicted perceptual bias from an optimal, reliability-weighted combination of the subject-and condition-specific predictions (Fig. 3a) and the subject-specific estimates from the control task (Fig. 2g). (**e**) Contrast values from a linear model describing individual subject (circles) and model (each diamond represents analyses based on the same sound sequence experienced by the subject connected by a line) perceptual bias in terms of (see inset in Fig. 3e): 1) the difference between change-point and non-change-point trials (CP), 2,3) the linear trend from SAC 2–6 for low- (Exp_low_) or high- (Exp_high_) noise trials, and 4) the difference between the two noise conditions (Noise). Asterisks indicate sign-rank test for *H_0_*: median value from the subject data=0, *p*<0.05.

### Individual differences in the modulation of perceptual biases

The above analyses demonstrated that for individual subjects, dynamic changes in the relevance and reliability of priors within an experimental session were associated with changes in the degree to which those priors biased perception. We identified similar effects across subjects, implying that individual differences in perception can reflect differences in how priors are updated and maintained in dynamic environments. Specifically, we compared subjects’ overall biases to the variability of their sensory and prediction errors (linear regression of the mean perceptual biases of individual subjects from non-change-point trials as a function of the STD of perceptual errors from the control task and the STD of prediction errors across non-change-point trials from the dynamic task; *F* statistic=7.39, *p*=0.002). According to these fits and consistent with Bayesian theory, subjects with higher overall prior-driven perceptual biases tended to have higher sensory variability (*β*=0.033, *t*-test for *H0*: β*=0, p*=0.013; Fig. 5a) and lower prediction variability (*β*=-0.030, *p*=0.002; Fig 5b). We also found individual differences in how perceptual biases changed as a function of particular task conditions, and that those differences were predicted by subject-specific changes in priors under those conditions. Subjects whose priors improved (i.e., became less variable) the most also tended to have the largest increases in prior-driven perceptual biases: 1) just after a change-point (Fig. 5e), 2) on experiencing samples from a new distribution (in the low- but not high-noise condition; Figs. 5c and d), or 3) between the high- and low-noise conditions (Fig. 5f). Thus, on average, subjects tended to weigh prior and sensory information according to their relative reliabilities, taking into account variability in the priors across task conditions and individual subjects.

**Figure 5:**
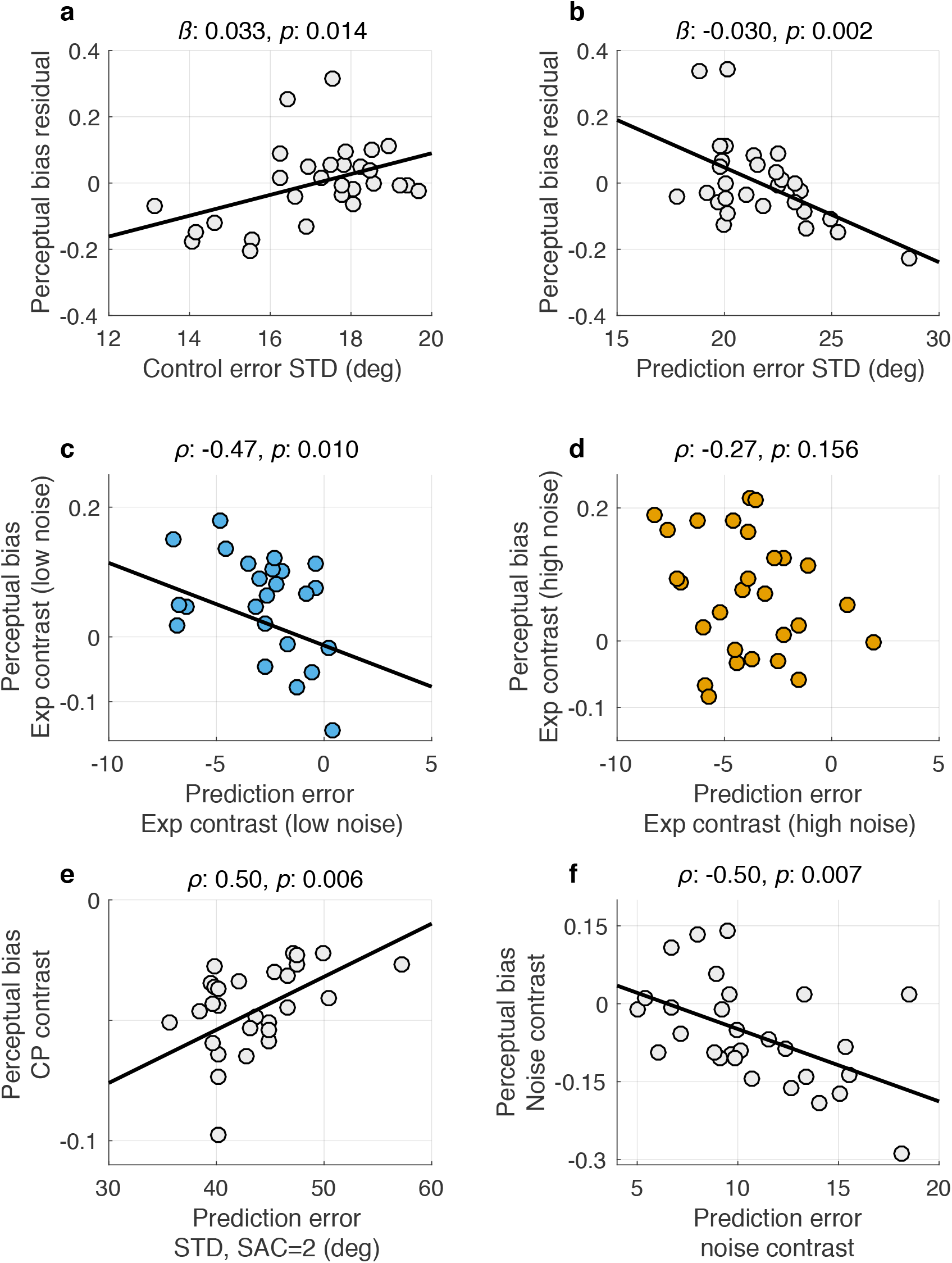
Individual differences in perceptual bias. (**a, b**) Relationship between overall (mean) perceptual bias and either overall localization ability (STD of perceptual errors on the control task, **a**) or overall prediction ability (STD of prediction errors from non-change-point trials on the dynamic task, **b**), after accounting for the other factor (hence “residual”) via linear regression. (**c**–**f**) The dependence of perceptual bias on various task conditions, plotted as functions of the dependence of prediction-error STD on the same conditions: **c, d**) the linear trend from SAC 2–6 in the low-noise (**c**) and high-noise (**d**) condition (Exp); **e**) change-point versus non-change-point trials (CP); and **f**) high-versus low-noise condition (Noise). Points in each panel represent data from individual subjects. Lines are linear regressions.

To more quantitatively account for the factors that affected perceptual biases across task conditions and individual subjects, we used a linear model that included normative and non-normative terms that each were weighed according to their contributions to each subject’s behavior (Fig. 6). The normative terms were extracted from a Bayesian model of perception, which generated perceptual biases that minimized simulated perceptual errors, given each subject’s variable predictions and sensory estimates. These terms were: 1) prior relevance, which reflected the probability that the current sound came from the same generative distribution as the previous sound (and thus is related to the CP effects illustrated in Figs. 3 and 4; Fig. 6c); and 2) prior reliability, which reflected changes in the total width of the predictive distribution relative to the likelihood, given new samples (and thus is related to the Exp and Noise effects illustrated in Figs. 3 and 4; Fig. 6d). The non-normative terms included one describing a fixed bias as a function of the prediction error, one to allow the strength of perceptual bias to depend on reported confidence (i.e., whether the subject bet high or not), and spatial terms to account for the subjects’ overall tendency to give perceptual reports that were biased slightly towards straight ahead (Fig. 2f). On average, the linear model captured the behavioral trends well (Fig. 6b), based on contributions of each of the terms described above that tended to vary in magnitude across subjects (Fig. 6e). By comparison, a parameter-free normative model captured some of the behavioral trends (Fig. 6a) but reported higher perceptual biases than subjects (compare red points and bar in Fig. 6e), particularly on change-points (compare green points and bars in Fig. 6e).

**Figure 6:**
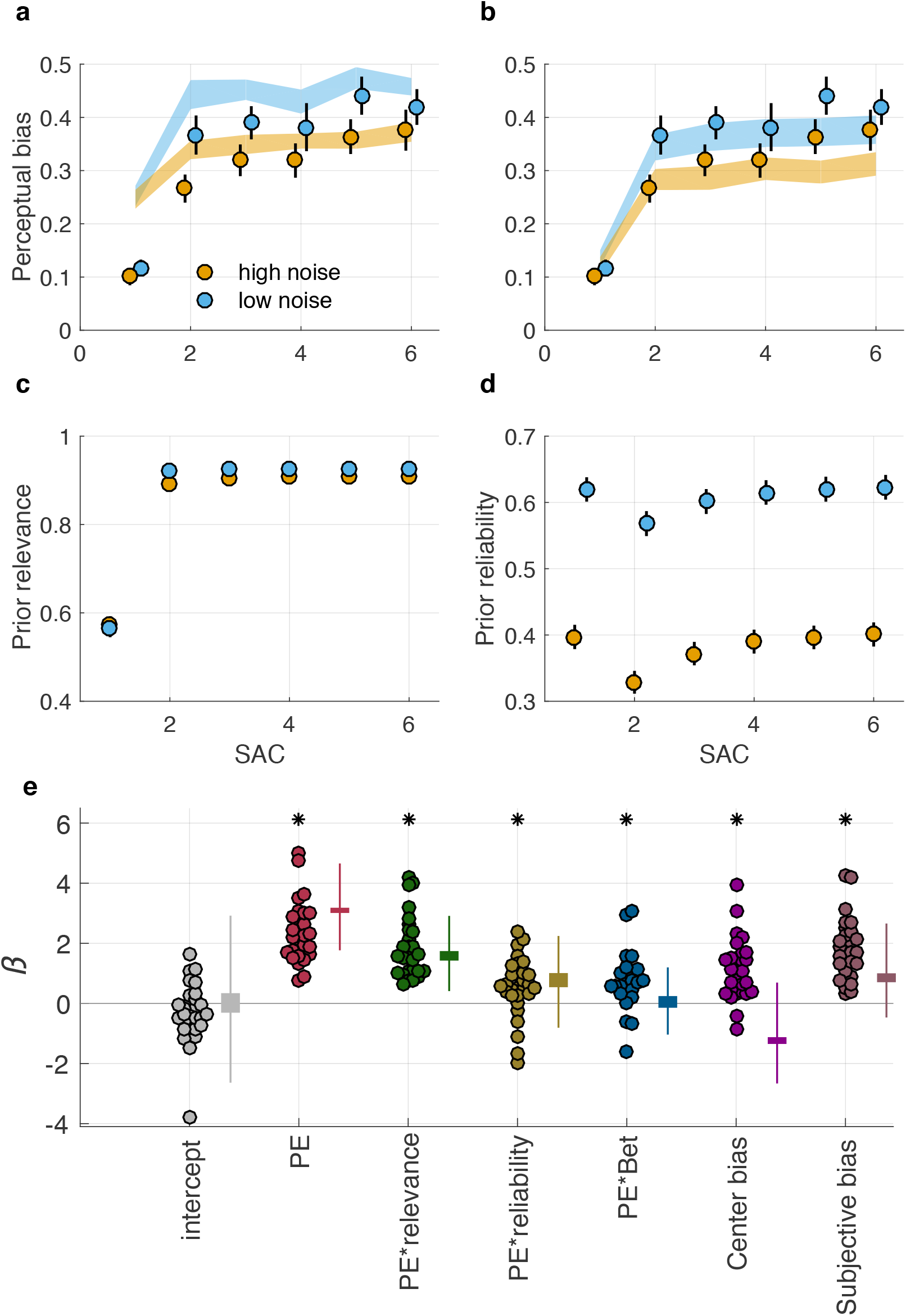
Dynamic modulation of perceptual bias by normative and non-normative factors. (**a**) Comparison of a parameter-free normative model (ribbons indicate mean±SEM simulated perceptual bias for the same task sequences experienced by the subjects) and the subjects’ behavior (points and errorbars are mean±SEM across subjects), shown as a function of sounds after a change-point (SAC) for the two noise conditions (colors, as indicated). (**b**) Comparison of the linear model shown in panel **e** to behavior. Conventions as in panel **a**. (**c**,**d**) Dependence of the normative factors used in both models on task conditions: (**c**) prior relevance, which measures the probability of the current sound coming from the same distribution as the previous sound; and (**d**) prior reliability, which measures the anticipated precision of the predictive distribution relative to the likelihood distribution prior to stimulus presentation. (**e**) Best-fitting parameter estimates from the linear model fit to behavioral data from each subject (points) and to simulations of the parameter-free normative model (thick and thin bars indicate 95% confidence intervals over simulated subjective values and over simulated mean values across subjects, respectively). PE=prediction error. Asterisks indicate coefficients with mean values that differed from zero (t-test, *p*< 0.05).

### Modulations of perceptual biases reflected in pupil diameter

A key question addressed in this work is whether arousal systems, as reflected in pupil diameter, contribute to the dynamic modulation of perceptual biases. Using linear regression at each time-point relative to sound onset (the average sound-evoked pupil response from all probe trials and subjects is shown in Fig. 7a), we found that pupil diameter varied with several of the factors from the linear model that accounted for behavioral biases (Eq. 6; Fig. 7b,c). Specifically, prior reliability was reflected in the baseline diameter before presentation of the probe sound, with smaller baselines reflecting more reliable priors (*p*=0.03, brown point in Fig. 7b). However, prior reliability did not modulate the magnitude of the stimulus-evoked pupil response, after accounting for the baseline effect. In contrast, prior relevance was unrelated to baseline diameter but was robustly encoded by the stimulus-evoked pupil diameter, with larger evoked pupil responses reflecting lower prior relevance. This effect peaked around the time of the maximum sound-evoked pupil response (permutation test for effect duration: duration=1.0 s, *p*=0.02). The pupil response also reflected the subjects’ upcoming bet, with high bets corresponding to larger pupil diameters, particularly late in the fixation interval (duration=1.8 s, *p*=0.01).

**Figure 7:**
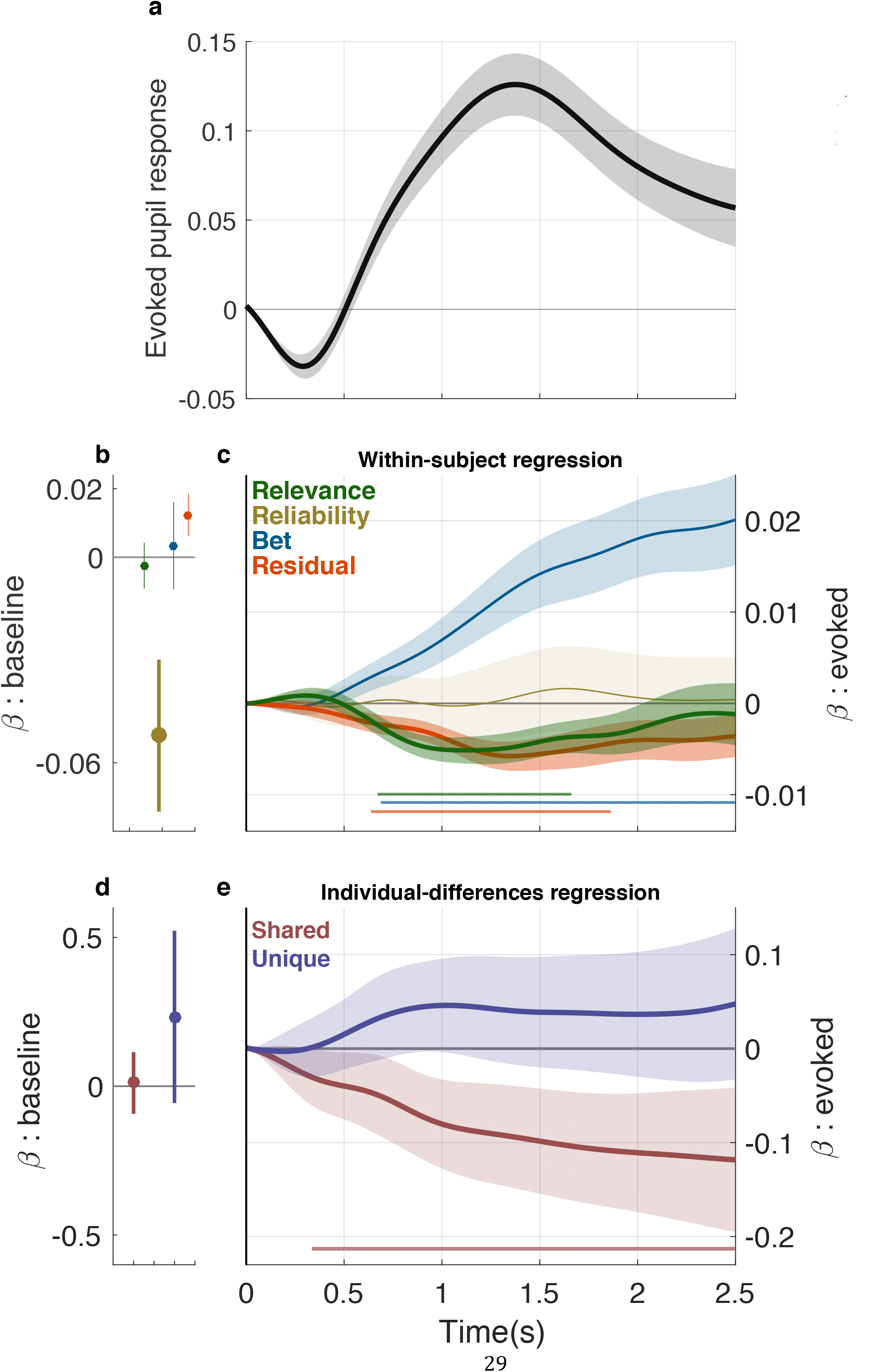
Pupil diameter reflects dynamic modulations of perceptual bias within and across individual subjects. (**a**) Mean±sem of per-subject mean evoked pupil response, defined as the pupil diameter relative to baseline during the measurement period. (**b,c**) Regression coefficients from a linear model accounting for modulation of baseline pupil diameter (**b**) or the evoked response (**c**) at each time-point using as predictors: 1) prior relevance, 2) prior reliability, 3) the upcoming bet, and 4) the residual perceptual bias from the linear model in Fig. 6d. Points and error bars in **b** and lines and ribbons in **c** represent mean±sem of values computed per subject and thus represent within-subject modulations. (**d,e**) Regression coefficients describing the relationship between shared or unique variance (colors, as indicated) in PE and PE*relevance coefficients from the behavioral model and average baseline (**d**) or stimulus evoked (**e**) pupil diameter. Points and error bars in **d** and lines and ribbons in **e** represent the correlation coefficient and 95% confidence intervals of the estimate and thus represent across-subject modulations. Abscissa in **a**, **c**, and **e** represents time relative to stimulus onset. Bold symbols in **b**, **d** or horizontal lines in **c**, **e** indicate periods for which *H*0: value=0, *p*<0.05 after accounting for multiple comparisons.

If the arousal system is contributing to the dynamic regulation of the influence of priors on perception, then pupil diameter may co-vary with adjustments in prior influence even after accounting for all of the factors in the behavioral linear model (for example, if variability in internal representations of sound-source location affect both behavior and arousal). We therefore included the residual perceptual bias from our model of behavior (Fig. 6) in our model of pupil diameter. A positive/negative value of the residual biases indicates that the subject was more/less biased by the prior on the given trial than predicted by the linear model. There was a trend toward positive coefficients for this term in explaining baseline pupil diameter (larger baseline diameters corresponded to slightly stronger biases than predicted by the behavioral model; *p*=0.06; Fig. 7b). In addition, there was a robust reflection of the residual bias term in sound-evoked pupil response (smaller responses near the peak of the evoked response corresponded to stronger biases than predicted by the behavioural model; duration=1.2 s, *p*=0.02; Fig. 7c). This residual bias effect implies that pupil diameter reflects not just particular factors like prior reliability and relevance that can be used to make effective predictions in dynamic environments (Nassar et al., 2012), but also the extent to which those and other factors are actually used to bias perception from one stimulus to the next.

In addition to these average, within-subject effects, there were also across-subject relationships between pupil diameter and perceptual biases, particularly with respect to prior relevance (Fig. 7d,e). We constructed a new linear model to account for these relationships. Because the behavioral influences of prior relevance (PE*relevance term in Fig. 6e) and overall perceptual biases (PE term in Fig. 6e) covaried considerably across subjects (*r*=0.77, *p*<10^−9^), we included in the pupil regression two individual-difference variables that corresponded to the shared and unique variance of the two behavioral coefficients. The effects of the shared term were negative for most of the measurement window (Fig. 7e; duration=2.2 s, *p*=0.01). In contrast, the unique-variance term did not show a strong relationship to average pupil traces. This result implies that subjects who had the strongest overall perceptual biases, and modulated them most according to prior relevance, tended also to have the smallest stimulus-evoked pupil responses.

To further quantify these within- and across-subject relationships between pupil diameter and task performance, we used pupil diameter to predict the subjects’ perceptual biases. Specifically, we created three normalized variables to reflect within- and across-subject variability in pupil responses at the time of peak modulation for residual bias (2.1 s following stimulus onset) along with their multiplicative interaction. Each pupil-derived variable was included as a modulator of prediction errors in three different linear models of perceptual errors. In the simplest model, pupil-derived measures alone predicted systematic differences in perceptual biases observed in the behavioral data (Fig. 8a), such that biases were: 1) larger for trials in which pupil responses were smaller than average (*t*-test, *p*<10^−4^), 2) larger for subjects who had smaller than average pupil responses (*p*<10^−3^), and 3) modulated from trial to trial more steeply for subjects with smaller overall pupil responses (*p*<0.01; Fig. 8b). Consistent with these relationships, the pupil-based measures offered a substantial improvement to the base model in terms of predicting behavior (likelihood-ratio test, *p*<10^−7^; Fig. 8c). The pupil-based measures also offered an explanatory advantage when added to more complex models that accounted for direct fixed effects (one coefficient for all subjects) or random effects (one coefficient per subject) of stability, reliability, and bets on perceptual biases (*p*<10^−4^ for both models; Fig. 8c). Taken together, these results imply that fluctuations in pupil diameter, particularly those mediated by stimuli and related to context stability, can be used to determine the extent to which perception is biased towards pre-existing priors.

**Figure 8:**
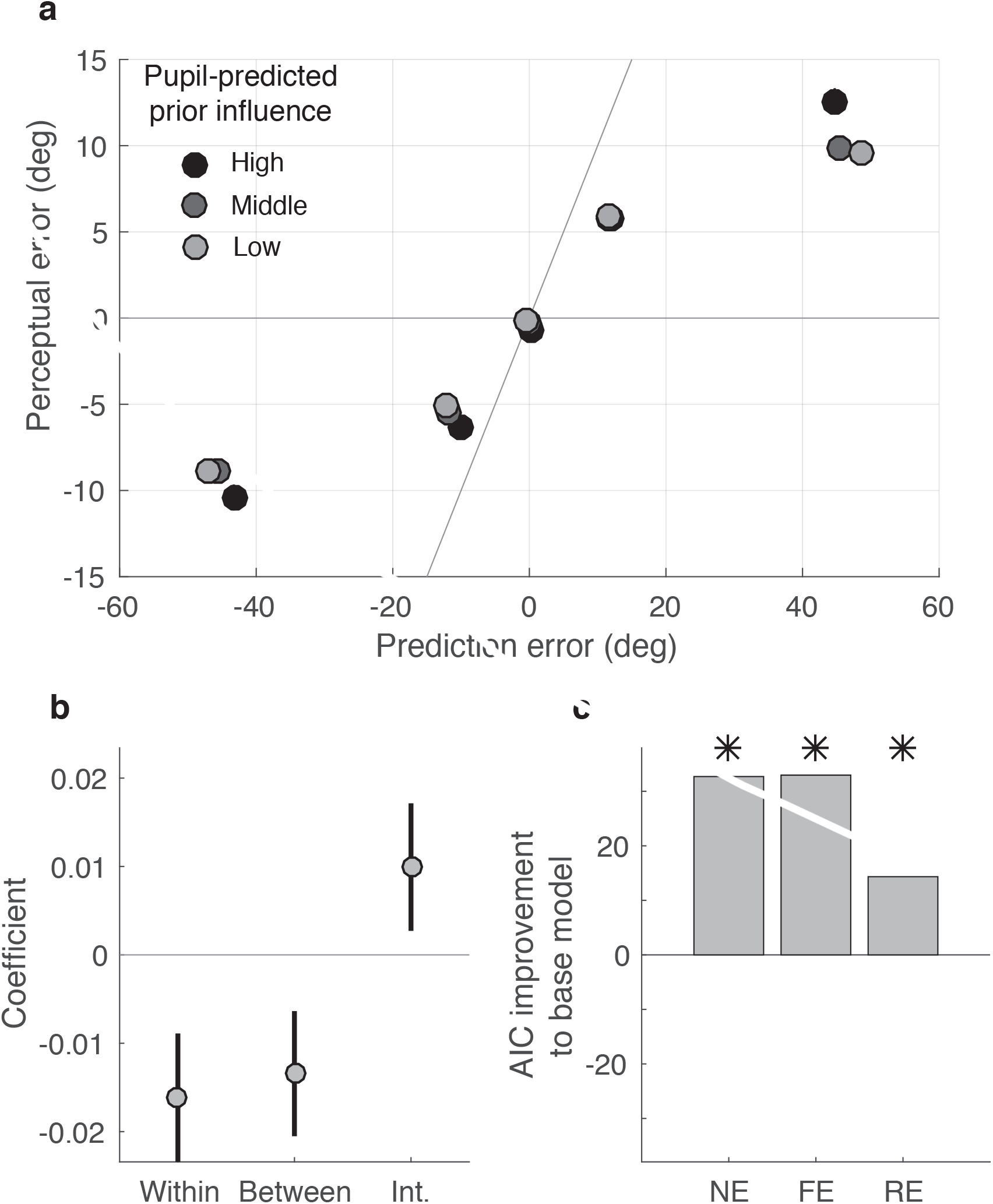
Pupil diameter predicts perceptual bias. (**a**) Perceptual error, sorted according to the pupil-predicted prior influence (gray scale, as indicated, corresponding to the bottom quartile, middle 50%, and top quartile) and plotted according to prediction error. Points are mean values computed across all subjects. (**b**) Mean±95% confidence intervals for pupil coefficients describing within- and between-subject effects of pupil diameter, as well as their interaction. See text for details. (**c**) Improvement in AIC achieved by adding pupil-based predictors to models that include: 1) a fixed perceptual bias across all subjects (NE), 2) a fixed perceptual bias and fixed model-based effects of perceptual bias across all subjects (FE), and 3) a random effects model that fits all bias and modulation terms separately for each subject (RE, which is equivalent to the normative linear model in Fig 6). Asterisks indicate significant improvements (likelihood-ratio test, *p*<0.05).

## Discussion

We used a novel auditory-localization task to show that the influence of prior expectations on perception is regulated rapidly and adaptively in changing environments. This work combines and extends several lines of research. The first has emphasized the role of priors on the perception of an uncertain sensory stimulus (Knill and Richards, 1996). Many of these studies have focused on priors that are related to relatively stable properties of the environment (W. J. Adams et al., 2004; Stocker and Simoncelli, 2006), although several recent studies have shown that certain sensory or sensory-motor priors can be learned relatively rapidly (W. J. Adams et al., 2004; Berniker et al., 2010; Burge et al., 2008; Stocker and Simoncelli, 2006; Tassinari et al., 2006). The second has shown that under a variety of conditions, individual variability in decision-making can involve differential use of priors (Stanovich and West, 2000). The third has identified how predictions are updated and used to make decisions in dynamic environments (Behrens et al., 2007; Nassar et al., 2010; Wilson et al., 2013). The fourth has related this dynamic updating process to changes in physiological arousal (Nassar et al., 2012; Preuschoff, 2011). We showed that many of these principles, including dynamic, arousal-related adjustments in predictions, apply to how priors are updated and used to guide perception.

These principles involve ongoing assessments of the relevance and reliability of priors that represent a form of statistical learning (Tenenbaum et al., 2011; Vapnik, 2013). We quantified this learning process using two variables derived from normative theory (R. P. Adams and MacKay, 2007; Mathys, 2011; Nassar et al., 2010; O'Reilly, 2013; Payzan-LeNestour and Bossaerts, 2011; Preuschoff and Bossaerts, 2007; Yu and Dayan, 2005). The first, which we termed prior relevance, is closely related to unexpected uncertainty and reflects the probability that a new observation is consistent with recent history (Nassar et al., 2010; Yu and Dayan, 2005). The second, which we termed prior reliability, is a form of reducible uncertainty that reflects ambiguity, typically resulting from undersampling, about the current generative process (Payzan-LeNestour and Bossaerts, 2011; Preuschoff and Bossaerts, 2007). Previous work showed that new information exerts the least influence on existing predictions when those predictions are the most relevant and reliable (Behrens et al., 2007; Nassar et al., 2010). We showed analogous effects for perception: new sensory input exerts the least influence on perception, relative to the influence of priors (i.e., perceptual biases are largest), when the priors are the most relevant and reliable.

Both of these normative factors, scaled according to their effects on each subject’s behavior, were reflected in modulations of arousal state as measured by pupil diameter. Prior reliability corresponded to changes in baseline pupil diameter, and prior relevance corresponded to changes in the stimulus-evoked change in pupil diameter. These modulations were similar to our previous findings in a predictive-inference task (Nassar et al., 2012). Here prior relevance was the stronger effect, with larger, stimulus-evoked pupil dilations reflecting the relatively diminished effects of priors on perception just after a change-point. This result is consistent with the idea that transient increases in arousal, in response to surprising events or other factors, may generally correspond to high sensitivity to immediate sensory input, possibly from a reduced influence of prior expectations (Pfaff, 2006; Sara and Bouret, 2012). This result encompassed both within- and across-subject effects, suggesting that surprise-related modulations of arousal may be an important source of individual variability in perceptual abilities.

We also found relationships between subjective confidence, perceptual biases, and pupil diameter. We measured confidence using a post-decision bet, which previously has been linked to the sensory-driven decision variable that also governs the speed and accuracy of the perceptual decision (Kepecs et al., 2008; Kiani and Shadlen, 2009; Persaud et al., 2007). We showed that confidence is also modulated according to changes in the relevance and reliability of perceptual priors that affect perceptual errors. This modulation was also evident in pupil diameter, which reflected bet frequency even after accounting for the normative variables that also governed the perceptual biases. However, this extra effect was in the opposite direction as for the normative factors, relative to the behavioral effect: high bet frequency corresponded to larger pupil diameters but stronger prior influence. This pupil effect is somewhat surprising given that pupil diameter can be enhanced under uncertain, rather than certain, conditions (Jepma and Nieuwenhuis, 2011; Lempert et al., 2015; Nassar et al., 2012; Satterthwaite et al., 2007; Wessel et al., 2011) (but see (Preuschoff, 2011)). One possible explanation for this discrepancy is that the post-decision bet led subjects to anticipate the increased reward or risk associated with high-bet trials, leading to stronger arousal responses (Manohar and Husain, 2015; Satterthwaite et al., 2007). This idea is supported by the time course of bet-related pupil dilations, which had a maximal dilation immediately prior to the perceptual report. This idea also highlights the multiple, possibly interacting roles that the arousal system likely plays in even simple sensory-motor tasks like this one.

These multiple roles undoubtedly result from multiple mechanisms by which arousal affects neural information processing (Robbins and Everitt, 1995). One such mechanism likely involves cortical levels of norepinephrine (NE), which is controlled via neurons in the midbrain nucleus locus coeruleus (LC) (Aston-Jones and Cohen, 2005). Firing rates of LC neurons correlate with pupil diameter over relatively short timescales, which has prompted the suggestion that pupil diameter can be used as a proxy for LC activity (Aston-Jones and Cohen, 2005; Bouret and Richmond, 2015; Joshi et al., 2016; Nieuwenhuis et al., 2011). Thus, the factors in our task that corresponded to larger pupil diameters, such as more surprising stimuli with lower prior relevance, may also correspond to increased LC activation. This activation, in turn, would increase levels of cortical NE, which have been theorized to signal unexpected context changes and allow neural representations to reorient rapidly to meet changing contextual demands, possibly via modulations of the input/output gain of individual cortical neurons (Aston-Jones and Cohen, 2005; Bouret and Sara, 2005; *MatherClewettSakakiHarleyinpress*, 2015; Servan-Schreiber et al., 1990; Yu and Dayan, 2005). In principle, such a mechanism seems well poised to regulate the influence of prior expectations on sensory input. Future work should test this idea directly.

## Experimental Procedures

Human subject protocols were approved by the University of Pennsylvania Internal Review Board. 29 subjects (16 female, 13 male) participated in the study after providing informed consent.

### Auditory-localization task

We used a novel auditory-localization task in which subjects heard sounds with varying source locations that were simulated using head-related transfer functions (HRTFs) from the IRCAM database (http://recherche.ircam.fr/equipes/salles/listen/download.html). Each sound was a sequence of five Gaussian noise pulses bandpass filtered between 100 Hz and 15 KHz. The pulses were 50 ms each with a delay of 10 ms between each pulse. For each subject, we tested a number of HRTFs during the initial session by playing sound sequences that circularly traversed the entire horizontal plane in 15° intervals. We picked the HRTF that was reported to give the most uniformly circular percept for the sound sequence. Each subject performed 3–6 total sessions.

Each subject completed two tasks per session. The first was a control localization task that required the subject to indicate the perceived location of simulated sound sources that were sampled independently and uniformly randomly along the frontal, horizontal plane. In each of 72 trials, the subject was required to: 1) fixate for 2.5 s while listening to the auditory stimulus; and 2) indicate the perceived location of the sound using a mouse, which controlled a cursor that moved along a semi-circular arc on the computer screen that represented the range of possible sound-source locations (Fig. 1). Failure to maintain fixation resulted in a warning sound and trial break. Feedback was displayed on the screen after the subject reported the perceived location.

The second task was a dynamic localization task that required the subject to report predictions, perceptions, and confidence judgments of sound-source locations that were generated from a change-point process along the same horizontal plane. For this task, the subject listened to extended sequences of auditory stimuli generated by the change-point process, paired with visual cues indicating the presented source location on the semi-circular arc. During the presentation of these sequences, no action was required. Occasionally, however, the sequences stopped, indicating the start of a “probe trial” with the following structure (Fig. 1c). First, the subject was required to predict the angular location of the next, upcoming probe stimulus on the arc using a mouse. Second, following the prediction, the subject was required to maintain fixation for 2.5 s. The auditory probe stimulus, with no corresponding visual cue, was presented at the beginning of this fixation period. Fourth, after the fixation period ended, the subject indicated the perceived location of the probe stimulus using the mouse and the visual display. Fifth, the subject then bet on the accuracy of the perceptual report (Fig 1). Each subject performed four blocks of the dynamic task per session, which included ~30 probe trials each. Each session lasted ~90 min.

The sequence of simulated sound-source locations for the dynamic task was determined according to a process that included both irreducible variability (noise) and abrupt discontinuities (change-points). Source locations were sampled from a Gaussian distribution with a standard deviation (STD) that was held constant within a block of 600 trials (10° or 20° for the low- or high-noise condition, respectively) and a mean that underwent abrupt change-points with a fixed probability, or hazard rate (*H*), of 0.15 for each sound sample. At each change-point, the mean of the generative distribution was resampled uniformly across the sound space, such that the newly generated source locations were independent from previous ones. The sequence was interrupted for probe trials at random using a procedure that ensured roughly even distribution of probes occurring 1–6 sounds after a change-point (SAC). Thus, on some trials the probe sound-source location was independent of the previous stimulus sequence (SAC=1). On other trials, the probe location was generated from the same distribution that produced the immediately preceding locations (SAC=2–6).

Subjects were motivated to make accurate perceptual reports on each probe trial through an incentive system. They were instructed to bet high if they were confident that the true location was within a 16° window centered on their second (perceptual) report, and to bet low otherwise. A correct/incorrect high bet resulted in a score of (15/-10), and a correct/incorrect low bet resulted in a score of (5/-3). Subjects were paid a bonus that depended on their total score.

### Behavioral data analysis: contrasts

Probe trials were sorted into twelve conditions according to the recency of the previous change point (SAC=1–6) and noise condition (high/low). Perceptual and prediction errors were computed by subtracting reported percepts and predictions from the true (simulated) sound source location for each trial. For each condition, the STD of prediction and estimation errors was used as a metric of average absolute error magnitude.

To quantify how prediction errors, estimation errors, and average betting depended on specific task conditions, we performed four orthogonal linear contrasts. Each contrast was computed by multiplying a weight matrix by the measured prediction errors, estimation errors, or average betting, aggregated according to the twelve task conditions for a single subject. Weight matrices were mean centered and chosen to identify: 1) differences between change-point and non-change-point trials (CP); 2) linear increases with increases in the number of sounds experienced following a change-point, from SAC=2 to SAC=5, in the high-noise condition (Exp_high_); 3) comparable linear increases in the low-noise condition (Exp_low_); and 4) differences between the high- and low-noise conditions (Noise). Thus, the contrasts provided a per-subject measure of how much each behavioral measurement was modified according to these factors. For Figs. 3–5, we considered only sound sequences following relatively large change-points, corresponding to at least twice the generative STD.

### Behavioral data analysis: perceptual bias

To quantify the influence of the prior on the perceptual report, we measured the slope of the best-fit line to the relationship between prediction errors (prediction–true location) and perceptual errors (percept–true location) for the given task condition. Slopes close to one indicate a high perceptual bias, and slopes close to zero indicate low perceptual bias. To measure how the perceptual bias evolved as a function of the number of sounds after a change-point (SAC), we used the following linear model and included only data from sequences following noticeable change-points (jumps of at least twice the generative STD):

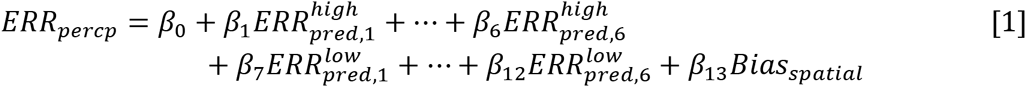

where *ERR_percp_* is the perceptual error and *ERR_pered,1_^high^* is the prediction error on change-point trials (SAC=1) in the high-noise condition, and so on. The last term, *Bias_spatial_,* captures the slight bias in the perceptual reports towards center of the screen.

### Behavioral data analysis: theoretical benchmarks

The theoretically expected overall perceptual-error STD per subject (abscissa in Fig. 2i) was computed from an optimal, reliability-weighted combination of prior and sensory information: 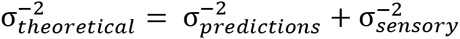. The theoretically expected perceptual-error STD per subject (diamonds in Fig. 3c,d), given their corresponding predictions for each SAC condition, was computed using 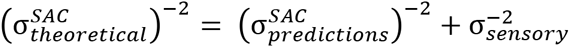. theoretically expected betting frequency (diamonds in Fig. 3e,f) was computed as the probability mass contained in a 16° window centered at the mean of a Gaussian with a STD of the theoretically expected perceptual errors, 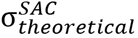. Thus, the proportion of expected high bets increased with narrower perceptual error distributions. The theoretically expected perceptual bias per subject (diamonds in Fig. 4d,e) was computed as 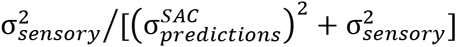. In all of the above, 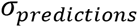 is the STD of prediction errors on non-change-point trials, 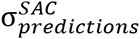 is the STD of prediction errors for the specified number of sounds after a change-point, and 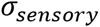 is the STD of perceptual errors on the control task, computed per subject.

### Behavioral data analysis: normative model

Auditory localization in a dynamic environment can be posed as a perceptual inference problem with the goal of inferring the location of the sound source on trial (*X_t_*) according to a noisy internal sensory representation of that sound source (*λ_t_*) and the history of previously experienced sound sources (*X_1:t-1_*). This problem can be simplified by exploiting the conditional independencies in the Markov change-point process through which sound sources are selected (see Supplementary Fig. 1). In particular, the sound sources locations on the current trial (*X_t_*) are independent of those on previous trials (*X_1:t-1_*) conditioned on the mean of the generative distribution on the current trial (μ_t_). In turn, the mean of the generative distribution on the current trial (μ_t_) is also independent of previous observations conditioned on: 1) the mean of the generative distribution on the previous trial (μ_t-1_), and 2) a latent change-point variable that determines whether the mean is resampled on the current trial (s_t-1_). Applying Bayes rule to invert the generative graph in Supplementary Fig. 1 yields the following inference equation:

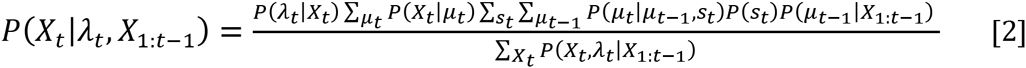

where the likelihood P(λ_t_|X_t_) reflects the conditional distribution of internal representations across true sound source locations; P(λ_t_|μ_t_) reflects the conditional probability of a sound source location being generated from a particular generative mean; 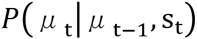 reflects the process through which means are resampled on change-point (s_t_=1) trials; and P(s_t_) is the hazard rate (*H*), which was fixed to 0.15 for the task and all simulations. The likelihood P(λ_t_|X_t_) was modeled as a normal distribution centered on X_t_ with a variance that was fixed for each subject to the variance of perceptual reports made by that subject on the control task 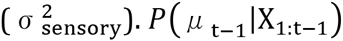 is the distribution over possible generative means, which can be updated recursively. Although exact Bayesian solutions to this recursive problem exist (R. P. Adams and MacKay, 2007; Wilson et al., 2010), we use a normal approximation to the Bayesian mixture distribution with a mean 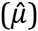1 and variance 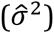 that capture the key dynamics of normative inference and offers better descriptions of human behavior (Nassar et al., 2010). As in previous work, predictions made using this approximation were more accurate than subject predictions. To account for this discrepancy, we created a subjective prediction 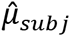 by sampling a random normal variable with mean equal to 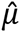 and a variance that was incremented on each trial according to the difference in variance of subject and normative prediction errors:

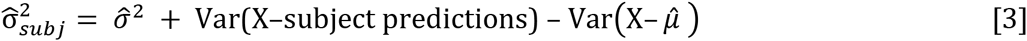

Thus, the overall model incorporates the prospect of sub-optimal predictions about the sound-source location but implements Bayesian optimal combination of these predictions with incoming sensory information according to environmental dynamics.

Perceptions and predictions from the normative model were simulated by sampling internal representations (λ_t_) and subjective predictions 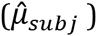 for each trial according to the actual sequence of sound source locations experienced. Descriptive statistics for model simulations were averaged across 100 such simulated runs.

In addition to simulating behavioral data, the normative model also allowed us to extract latent variables that guide normative adjustments in perceptual bias. In particular, the model adjusts bias towards prior expectations in accordance with the relevance and reliability of those expectations. The relevance of prior expectations (π_t_) is, in our generative framework, equal to the probability that a change point did not occur on this trial given all previous data. This probability was computed on each trial by marginalizing Eq. 2 over all dimensions other than *s*. The impact of normative priors also depends critically on their reliability relative to that of the likelihood distribution capturing the noisy internal sensory representation (λ_t_):

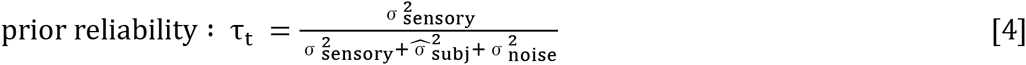

where τ_t_ is prior reliability, 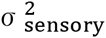 is the variance of perceptual reports made by that subject on the control task, 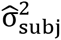 is the variance on subjective assessments of the underlying mean, and 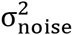 is the expected variance of sound source locations about that mean. The sum of the latter two terms reflects the total variance on the model’s predictive distribution over possible sound locations. To ensure that these latent variables best reflected circumstances experienced by the subject, we fixed the model predictions 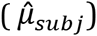 to the actual subject predictions from each trial and computed each measure as the expected value across all possible values of λ_t_ using a grid-point approximation.

### Behavioral data analysis: Linear model of perceptual bias

To measure how the prior bias was modulated by normative and other factors, we fit the following linear model to data from all change-point and non-change-point trials:

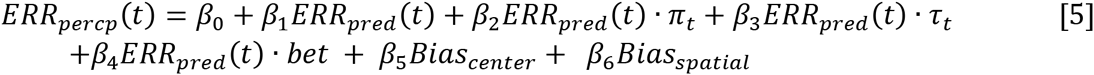

where β_1_ describes the overall prior bias; β_2_ and β_3_ describe the extent to which the overall bias is dynamically modulated by the prior relevance and reliability, respectively (see above); β_4_ describes the interaction of prior bias with betting (a binary variable); β_5_ describes the bias towards the center of the screen; and β_6_ captures the angular spatial bias (mean perceptual error at the given angle) measured in the control task. Residuals from the model fit were signed according to the prediction error on each trial to create a residual bias term for use in pupil analysis.

### Pupil measurements

Pupil diameter was sampled from both eyes at 60 Hz using an infrared eye-tracker built into the monitor (Tobii T60-XL). Pupil analyses used the mean value from the two eyes at each time point measured during fixation. We excluded from further analyses trials with blinks, which we identified using a custom blink-detection routine on the basis of pupil diameter and vertical and horizontal eye position. The raw pupil diameter was low-pass filtered using a Butterworth filter with a cut-off frequency of 10 Hz. These filtered measurements were then z-scored in each session. We also removed a linear trend in the average pupil diameter over the whole session to account for any slow drift. The pupil baseline for each probe trial was defined as the mean of the first three samples immediately preceding the measurement period for that probe trial.

### Linear model relating pupil diameter to behavioral parameters

To measure how the variables driving behavior were encoded in pupil diameter, we used the following linear model to explain the fluctuations in: 1) the baseline pupil diameter, and 2) stimulus-evoked pupil response across the 2.5 s following the auditory stimulus:

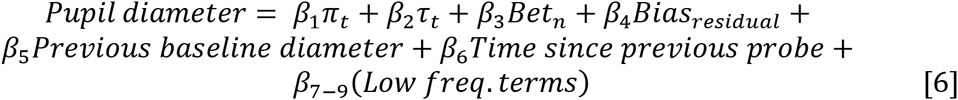

where τ_t_ and π_t_ are the reliability and relevance, respectively; β1-4 capture relationships between pupil diameter and the computational and behavioral variables of interest; β5-6 capture persisting fluctuations in pupil diameter that are attributable to the previous trial; and β7-9 includes a set of three low-frequency cosine components for each session that capture variability due to slow modulations or session based differences in pupil diameter. The exact form of the cosine components was (cos(π·k(2n-1)/2n), where *k=0,1,2*; *n* is the trial number within the session; and *N* is the total number of trials in the session. When this model was fit to evoked pupil responses, an additional nuisance variable was added to the explanatory matrix that accounted for trial-by-trial differences in baseline diameter.

Significance testing for evoked pupil coefficients was done through cluster-based permutation testing to account for multiple comparisons over time. In short, *t*-tests were performed on each set of coefficient values across subjects separately for each time point. Cluster size was determined according to the number of contiguous time points for which this *t*-test yielded *p*<0.05. Cluster corrected *p*-values were determined by comparing cluster sizes attained in this way to those from a permutation distribution of maximal cluster sizes (Nichols and Holmes, 2002).

### Pupil-predicted perceptual bias

Trial-by-trial pupil measurements were extracted for the time of peak modulation for residual bias from the behavioral model, as measured by the absolute group *t*-statistic. Trial-by-trial measurements from each subject were regressed onto a set of nuisance variables that included explanatory variables *β*_5+_ from Eq. 6 to remove variance attributable to potentially confounding factors. For each time point, residual pupil measurements were concatenated across subjects and then divided into two separate variables: one variable accounted for average measurements for each subject and one that reflected normalized deviations from the average measurement within each subject. The six resulting variable arrays were z-scored and multiplied by trial prediction errors to create a predictor matrix. Trial-by-trial perceptual errors were regressed onto three separate models with and without the inclusion of the pupil predictor matrix: 1) a base model including an intercept term and a prediction error term to capture fixed effects of perceptual bias across all subjects as well as the spatial bias terms described above; 2) a fixed-effects model that also included interaction terms accounting for modulation of perceptual bias by prior relevance and reliability the subjects’ betting; and 3) a random-effects model that included all terms in model 2 separately for each subject.

## Author Contributions

All authors designed the experiment and analyses and wrote the manuscript, KK collected the data, and KK and MRN analysed the data.

## Acknowledgements

We thank Ana-Andreea Stoica and Naim Kabir for aiding in data collection. We thank Long Ding and Takahiro Doi for helpful comments. This work was funded by NIH grants F32 MH102009 (MRN) and R01 EY015260 and NSF 1533623 (JIG).

---

**Supplementary Figure 1.**
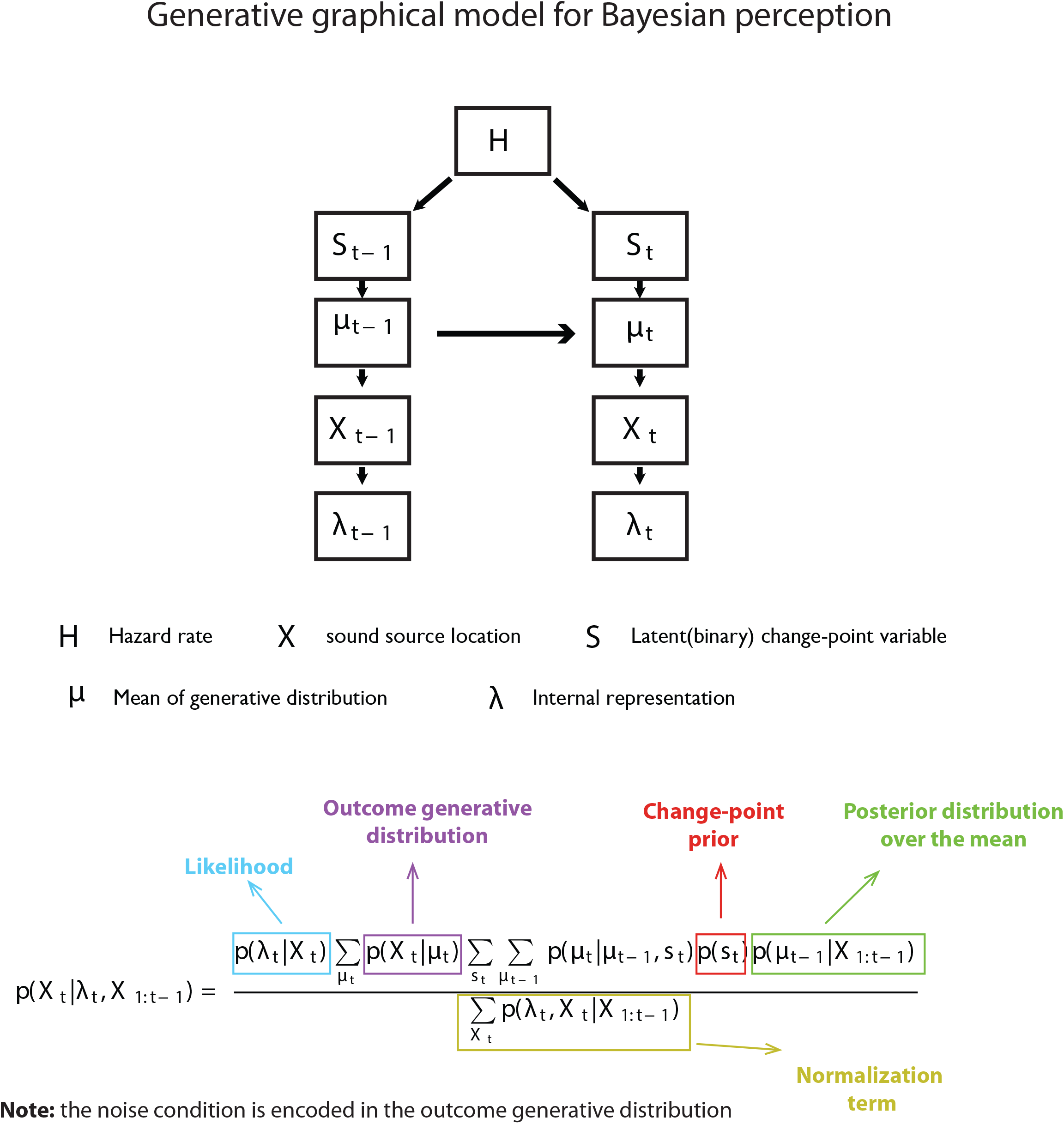

